# Identification, expression and subcellular localization of *Leishmania amazonensis* and *Leishmania infantum* Phospholipases A_1_

**DOI:** 10.64898/2026.03.27.714763

**Authors:** López Sebastián Andrés, DeSouza-Vieira Thiago Soares, Trinitario Sebastián Nicolás, Pereira-Dutra Filipe Santos, Rajão Matheus Andrade, Risso Marikena Guadalupe, Sanchez Alberti Andrés, Bivona Augusto Ernesto, Lauthier Juan José, Gimenez Guadalupe, Torres Bozza Patricia, Belaunzarán María Laura

## Abstract

Leishmaniases remain a significant global public health threat, with *Leishmania amazonensis* and *Leishmania infantum* representing the etiological agents of the cutaneous and visceral forms in the Americas, respectively. Building on our previous identification of Phospholipase A_1_ (PLA_1_) in *Leishmania braziliensis*, this study provides a comprehensive molecular, immunological, and biochemical characterization of PLA_1_ in *L. amazonensis* and *L. infantum* promastigotes. We analyzed PLA_1_ activity and expression, purified the recombinant enzyme from *L. amazonensis*, and validated protein expression using a specific anti-PLA_1_ serum. The major contribution of this research is the first description of the subcellular localization of a PLA_1_ within the *Leishmania* genus. Moreover, our results reveal an unprecedented association between PLA_1_ and lipid droplets within the parasites. This discovery is of particular interest as it provides the first evidence linking this enzyme to lipid storage organelles in *Leishmania*. Given that PLA_1_ is an established virulence factor in other trypanosomatids, these findings suggest a specialized role for the enzyme in parasite lipid metabolism and potentially in its pathogenic mechanisms, opening new perspectives for understanding *Leishmania* biology.

## 1. Introduction

Leishmaniases are caused by protozoan parasites from the genus *Leishmania* (Trypanosomatida: Trypanosomatidae), which are transmitted by female phlebotomine sandflies. These zoonotic and neglected tropical diseases comprise a public health problem, imposing a significant economic burden in the Americas [1]. It is estimated globally that more than 1 billion people live in endemic areas and are at risk of infection, leading to 50,000 to 90,000 new cases of visceral leishmaniasis (VL), and 600,000 to 1 million new cases of cutaneous leishmaniasis (CL) annually (World Health Organization, 2026). Leishmaniases present a broad spectrum of clinical manifestations, ranging from self-healing localized or multiple cutaneous lesions to mucosal lesions and the potentially fatal visceral form if untreated [2]. The infecting *Leishmania* species and the immune response triggered in the host determine severity and clinical manifestations [3]. Approximately 53 *Leishmania* species have been described, including five subgenera and complexes, being 31 species parasites of mammals and approximately 20 pathogenic to humans [4]. In Argentina, *Leishmania amazonensis* mainly causes CL [1], but the course of disease can differ in severity leading to either diffuse cutaneous leishmaniasis, which is associated to an absence of cell-mediated immune responses to the parasite [5] or disseminated cutaneous leishmaniasis, distinguished by multiple secondary lesions occurring at sites remote from the primary focus [6]. *Leishmania infantum* is the etiological agent of VL in the Americas. The complex interaction between the host immune response and *Leishmania* virulence factors is responsible for parasite persistence and dissemination. Recent works highlight the importance of parasite lipid droplets (LD), conserved cytoplasmic lipid-enriched organelles induced upon host interaction and inflammatory events, participating in cell signaling mainly in lipid metabolism and eicosanoid synthesis [7–9].

Phospholipases are ubiquitous and heterogeneous groups of enzymes that considerably vary in structure and function, being classified as A_1_, A_2_, C, and D according to the site of hydrolysis of phospholipids and lysophospholipids [10]. Phospholipases A (PLAs) are key regulators of cellular physiology and established virulence factors in a wide range of microorganisms. Phospholipase A_1_ (PLA_1_) exerts toxicity through phospholipid hydrolysis, generating lysophospholipids that disrupt host membrane integrity and intracellular signaling, thus contributing to cellular dysfunction, tissue damage, and disease progression [11,12]. Conversely, phospholipase A_2_ (PLA_2_) activity drives pro-inflammatory signaling by releasing arachidonic acid (AA), the precursor of eicosanoids. These mediators orchestrate immune responses; however, pathogens can exploit this pathway to modulate the host immune environment, favoring their own survival and persistence [13]. Regarding PLA_1_ activity in *Leishmania* spp., phosphatidylcholine PLA_1_ activity was reported in *Leishmania braziliensis* soluble fractions, showing two-fold higher levels in amastigotes than in promastigotes. This enzymatic activity was independent of divalent cations, with an optimum activity at pH 6.4. Further cloning and expression of the putative PLA_1_ gene resulted in an active recombinant PLA_1_ that displayed this activity, thus confirming the identity of the gene. Ortholog search for this gene in other species showed the presence of putative PLA_1_ genes throughout 12 species, including *L. amazonensis* and *L. infantum* [14]. PLA_2_ activity in *L. amazonensis* is associated with disease progression since the induction of PLA_2_ in parasites enhanced tissue injury and resulted in prostaglandin E2 (PGE2) generation, leading to immunosuppression and parasite persistence in the host [15]. Furthermore, bromoenol lactone, a specific PLA_2_ inhibitor, altered *L. amazonensis* viability, decreased both lesion size and skin parasitism in BALB/c mice, thus supporting the involvement of PLA_2_ in parasite physiology and virulence [16].

In the present study, we performed a molecular, immunological, and biochemical characterization of PLA_1_s in *L. amazonensis* and *L. infantum*. We assessed PLA_1_ activity and expression in *Leishmania* promastigotes, purified the recombinant enzyme, and generated a specific anti-PLA_1_ serum to validate protein expression. Furthermore, we showed that the subcellular localization of PLA_1_ is associated with lipid droplets within the parasites.

## 2. Materials and methods

### 2.1. Ethics statement

To perform this work, BALB/c mice were bred and maintained in the animal facility at the Instituto de Investigaciones en Microbiología y Parasitología Médica (IMPaM, Consejo Nacional de Investigaciones Científicas y Técnicas - Universidad de Buenos Aires), Buenos Aires, Argentina. All the procedures were approved by the Institutional Committee for the Care and Use of Laboratory Animals (CICUAL, Facultad de Medicina, Universidad de Buenos Aires, CD N°298/2019), in accordance with guidelines provided by the Argentine National Administration of Drugs, Food and Medical Devices (ANMAT), and the Argentinian National Service of Sanity and Agrifoods Quality (SENASA), based on the US NIH Guide for the Care and Use of Laboratory Animals. All methods were reported in accordance with the ARRIVE guidelines.

### 2.2. Parasites

*Leishmania (L.) amazonensis* (IFLA/BR/67/PH8 and MHOM/BR/75/Josefa), *Leishmania (V.) braziliensis* (MHOM/BR/75/M2904), and *Leishmania (L.) infantum* (MHOM/MA/67/ITMAP263) were used. Promastigotes were axenically grown in RPMI medium (Invitrogen, Grand Island, NY, USA) supplemented with 20 mg/L hemin and 20% fetal bovine serum (FBS) (Internegocios S.A., Buenos Aires, Argentina), at 26°C and maintained by weekly passages [14,17,18].

### 2.3. Parasite lysates

The parasites (1×10^8^/ml) were washed three times with phosphate-buffered saline (PBS). Next, they were suspended in the same buffer containing 1X protease inhibitor cocktail (Sigma Chemical Co., St. Louis, MO, USA) and finally disrupted by four freezing/thawing cycles. Following centrifugation of the samples (8,000 xg, 20 min, 4°C), the resultant supernatants, corresponding to the soluble fraction of the parasite lysis, were collected and stored at -80°C. Hereafter, these soluble fractions are referred to as parasite lysates and were used for PLA_1_ activity assays and immunoblot analyses [14].

### 2.4. Phospholipase assays

Phospholipase activity was measured in the different parasite lysates using 1-palmitoyl-2-{6-[(7-nitro-2-1,3-benzoxadiazol-4-yl) amino] hexanoyl}-sn-glycero-3-phosphocholine (NBD-PC) (Avanti Polar Lipids, Inc., Alabaster, AL, USA) as substrate [14]. Briefly, 50 μl of sample + 124 μl of 0.1 M Tris-HCl/0.1 M sodium citrate pH 4.8 + 6 μl of 300 μM NBD-PC plus 0.6 mg/assay of soybean phosphatidylcholine (PC) were incubated for 4 h or overnight at 37°C. Reaction was stopped by adding 0.9 vol of 0.2 M ammonia in methanol and 0.9 vol of chloroform. Fluorescence intensity was determined in the aqueous phase, as a direct measure of PLA_1_ activity, using a PerkinElmer LS 55 Fluorescence spectrometer (Waltham, MA, USA) with fluorescence excitation and emission wavelengths of 460 nm and 534 nm, respectively. Alternatively, the reaction was finished with chloroform/methanol (1:1 vol/vol), lipids extracted according to Bligh and Dyer (Bligh and Dyer, 1959), and separated by thin-layer chromatography (TLC) on Silica Gel 60 plates (Merck, Darmstadt, Germany) using chloroform/methanol/water (65:35:2.5, v/v). The generation of lysophosphatidylcholine (LPC) was quantified by densitometry using ImageJ v1.54m software [20]. PLA_1_ specific activity was expressed as nmoles of LPC released.min^-1^.mg^-1^ protein. To characterize *L. amazonensis* PLA_1_ activity in promastigote lysates, the fluorescent substrate NBD-PC was used to obtain the following parameters: (i) Optimum pH: PLA_1_ activity was assayed with a universal buffer 0.1 M Tris-HCl/0.1 M sodium citrate (pKa 3.4, 4.8, 6.4, 7.7), adjusted to different pH values (3.0–8.0), for 4 h at 37°C. (ii) Requirement of divalent cations: to analyze their effect on PLA_1_ activity, 2 mM CaCl_2_ or 2 mM MgCl_2_ were added to the assay at pH 4.8 and 37°C.

### 2.5. Recombinant L. amazonensis PLA_1_ expression and purification

The nucleotide sequence of a putative lipase (class 3) gene in *L. amazonensis* (LAMA_000646500) was previously obtained by ortholog search using the reference strain genome (MHOM/BR/73/M2269) available in the genomic resource TriTrypDB and the NCBI databases [14]. A signal peptide at the N-terminus of the amino acid sequence (positions 1–30, predicted with a 67% probability using SignalP v5.0 – DTU Health Tech) [21] was removed to facilitate recombinant expression in *Escherichia coli*. Therefore, *L. amazonensis* PLA_1_ partial gene sequence (GenBank ACCN OR545371) was synthesized into the pET-28a (+) expression vector (Novagen, Merck Millipore, Sweden) (pET28a(+)-rLamaPLA_1_) by GenScript Biotech Corporation (Piscataway, NJ, USA) with optimized codon composition for *E. coli*. Besides, the expression vector included His-tags at both the N- and C-terminal regions to facilitate affinity chromatography purification. The pET28a(+)-rLamaPLA_1_ was used to transform competent One Shot® BL21pLysS *E. coli* cells (Invitrogen, Carlsbad, CA, USA), and recombinant protein expression was induced in Luria Bertani media (LB) supplemented with 1 mM isopropyl-1-β-D-thiogalactopyranoside (IPTG) for 20 h at 24°C. The recombinantly expressed *L. amazonensis* PLA_1_ (rLamaPLA_1_), accumulated as inclusion bodies (IBs), was then purified under denaturing conditions (8M urea) using a nickel-nitrilotriacetic acid resin (Ni^++^-NTA) His-Pur™ (Thermo Fisher Scientific Inc.) and further dialyzed in 50mM Tris-HCl, 10% glycerol, pH 8 buffer for *in vitro* refolding. Aliquots were analyzed by SDS-PAGE followed by Coomassie blue staining to assess protein purity, and by immunoblotting with an anti-histidine (anti-His) antibody (Sigma Chemical Co.) to confirm protein identity, respectively [22].

### 2.6. Anti-rLamaPLA_1_ sera

An immunization protocol was performed to raise antibodies against rLamaPLA_1_. Female BALB/c mice (6-8 weeks of age, *n* = 6) were injected by the intraperitoneal route with 5 doses of 20 μg (130 μg/ml) of rLamaPLA_1_ + 2 mg of aluminum hydroxide–based adjuvant, administered at 21 day-intervals (days 1, 22, 43, 64, 85). Sera were collected, before starting and after finishing the immunization protocol (days 0 and 100, respectively), for the determination of specific immunoglobulin (Ig) titers by ELISA, and stored at -20°C until used [23].

#### 2.6.1. Specific anti-rLamaPLA_1_ Ig titer determination by ELISA

To determine total anti-rLamaPLA_1_-specific Ig titer by ELISA, MaxiSorp™ plates (Thermo Fisher Scientific Inc.) were coated with 0.5 μg of rLamaPLA_1_ as antigen in PBS (5 μg/ml) overnight at 4°C. Plates were then washed 3 times with 0.05% Tween-20-PBS (PBST) and blocked with 300 μl of a 3% bovine serum albumin solution in PBS (PBS-BSA) for 1 h at 37°C. The plates were washed 3 times with PBST and incubated for 1 h at 37°C with 100 μl of 3-fold serial dilutions of each serum in 1% PBS-BSA (1:100, 1:300, 1:900, 1:2700, 1:8100, and 1:24300, respectively). Plates were washed 5 times with PBST, incubated with 100 μl of the conjugate goat anti-mouse Ig-HRP (BD Pharmingen, San Diego, CA, USA) diluted to 1:10000 (v/v) in 1% PBS-BSA, for 1 h at 37°C, and after washing 5 times with PBST, they were incubated for 10 min with 50 μl of Tetramethylbenzidine (TMB, Thermo Fisher Scientific Inc.) and 50 μl of H_2_O_2_. The reaction was stopped with 100 μl of 2N H_2_SO_4,_ and the absorbance was measured at 450 nm in a microplate reader (RT-6000, Bio Rad). Titers were then calculated as the inverse of the highest dilution in which the optical density was higher than 0.1 [24].

### 2.7. PLA_1_ expression analyses by immunoblot

To analyze protein expression of native PLA_1_ in parasite lysates from different *Leishmania* species, immunoblot analyses were performed using anti-rLamaPLA_1_ serum. The different samples (∼1,5 μg protein/lane) were suspended in Laemmli sample buffer [25]+ 100 mM dithiothreitol and separated by 10% SDS-PAGE. Proteins were transferred to nitrocellulose membranes and blocked with PBS + 3% skimmed milk for 2 h at 37°C. After this, membranes were incubated with anti-rLamaPLA_1_ serum (1:1000) as primary antibody, overnight at 4 °C. The immunoblot was developed using the secondary antibody anti-mouse Ig-HRP (1:1000) (Santa Cruz Biotechnology, Santa Cruz, CA, USA) and the SuperSignal West Pico (Thermo Fisher Scientific Inc., Rockford, IL, USA) as chemiluminescent substrate, following the manufacturer’s instructions. Images were acquired using the UVP BioSpectrum® Imaging System (Analytik Jena AG, Jena, Germany). Ponceau S staining was used as a loading control. PLA_1_ expression levels were determined by densitometry using ImageJ v1.54m software [20], and the intensity of the signal of the PLA_1_ band was normalized to that corresponding to loading controls.

### 2.8. Effect of anti-rLamaPLA_1_ serum on PLA_1_ activity

To determine the effect of specific anti-rLamaPLA_1_ serum on the enzyme activity, 50 μl of *L. amazonensis* promastigote lysates (∼15 μg protein/assay) were pre-incubated with anti-rLamaPLA_1_ serum (1:1000) or normal mouse serum (1:1000) as a control, for 1 h at 37°C, and then PLA_1_ assays were performed as described above.

### 2.9. Immunolocalization of PLA_1_ and lipid droplets within Leishmania promastigotes

Briefly, *L. amazonensis* and *L. infantum* promastigotes (2×10^6^ parasites/ml) were fixed with 4% formalin onto coverslips and rehydrated with 0.22 μm-filtered water at room temperature (RT) for 5 min. Next, slides were blocked with 2% PBS-BSA and parasites permeabilized with 0.1% Triton X-100 + 0.2% PBS-BSA (PBS-TB) for 30 min. In order to localize PLA_1_, slides were incubated with anti-rLamaPLA_1_ serum (diluted to 1:20 or 1:40 in 0.2% PBS-BSA) overnight at 4°C followed by anti-mouse IgG - Alexa Fluor^TM^ 546 conjugate (diluted to 1:200 in 0.2% PBS-BSA, Invitrogen) in the dark for 1 h at RT. Lipid droplets (LD) were stained with BODIPY^TM^ 493/503 dye (1:5000, Invitrogen) for 10 min. Simultaneously, nuclei and kinetoplasts were identified using DAPI staining (diluted to 1:5000 in 0.2% PBS-BSA, Invitrogen) for 10 min. Finally, the coverslips were mounted on slides using an antifade mounting medium (VECTASHIELD®, Vector Laboratories Inc., USA). Fluorescence was analyzed by confocal microscopy (Laser scanning microscopy LSM710 Meta, Zeiss) [26].

### 2.11. Analyses of the subcellular localization of PLA_1_ by bioinformatics

To identify the subcellular localization of *Trypanosoma brucei* PLA_1_ (TbPLA_1_), the TrypTag resource was used [27]. Considering that this platform explores the localization of both N- and C-terminal fluorescently tagged *T. brucei* proteins, we searched for the corresponding homologous sequence of TbPLA_1_: Tb927.8.7440, belonging to the TREU927 reference strain. Hoechst 33342 was used for staining nuclei and kinetoplasts, and imaging was performed using a Leica DM5500 B epifluorescence microscope.

### 2.12. Statistical analyses

Results were expressed as mean ± SEM. Unpaired Student’s t-test was used to compare between groups, and analysis of variance (ANOVA) was used for multi-group analysis using GraphPad Prism 8 software (GraphPad Software, Inc., San Diego, CA). Statistically significant differences were represented as *p<0.05, **p<0.01; ***p<0.001 and ****p<0.0001.

## 3. Results

### 3.1. PLA_1_ expression and activity differ between L. amazonensis and L. infantum promastigotes

In previous studies, our laboratory has contributed to the knowledge of trypanosome PLA_1_, especially in *Trypanosoma cruzi* and *Leishmania braziliensis*. [14,28,29]. Herein, we extrapolate our experience to further investigate the PLAs in *L. amazonensis* and *L. infantum*. To this end, the fluorescent substrate NBD-PC was used to analyze the presence of these enzyme activities in *L. amazonensis* and *L. infantum* promastigote lysates. Fig. 1A shows the generation of lysophosphatidylcholine (LPC) (black arrows) as well as free fatty acids (FFA), thus indicating the presence of A_1_ and A_2_ activities in *L. amazonensis* promastigote lysates. In the case of *L. infantum*, no evident release of LPC or FFA was detected under our experimental conditions, as the observed signal was similar to the lane of the negative control (C). As a positive control, *L. braziliensis* promastigote lysate generated LPC and FFA, confirming the presence of both PLA_1_ and PLA_2_ activities. Considering that no PLA_1_ activity, but only PLA_2_ [16,30,31], has been described in *L. amazonensis*, we focused our analysis in the first one. Densitometric analyses of TLC plates determined that *L. amazonensis* PLA_1_ specific activity was approximately 2.3-fold higher than that of *L. braziliensis* (1.00 ± 0.05 and 0.51 ± 0.07 nmoles of LPC released.min_-1_.mg^-1^ protein, respectively) (Fig. 1B).

**Fig. 1.**
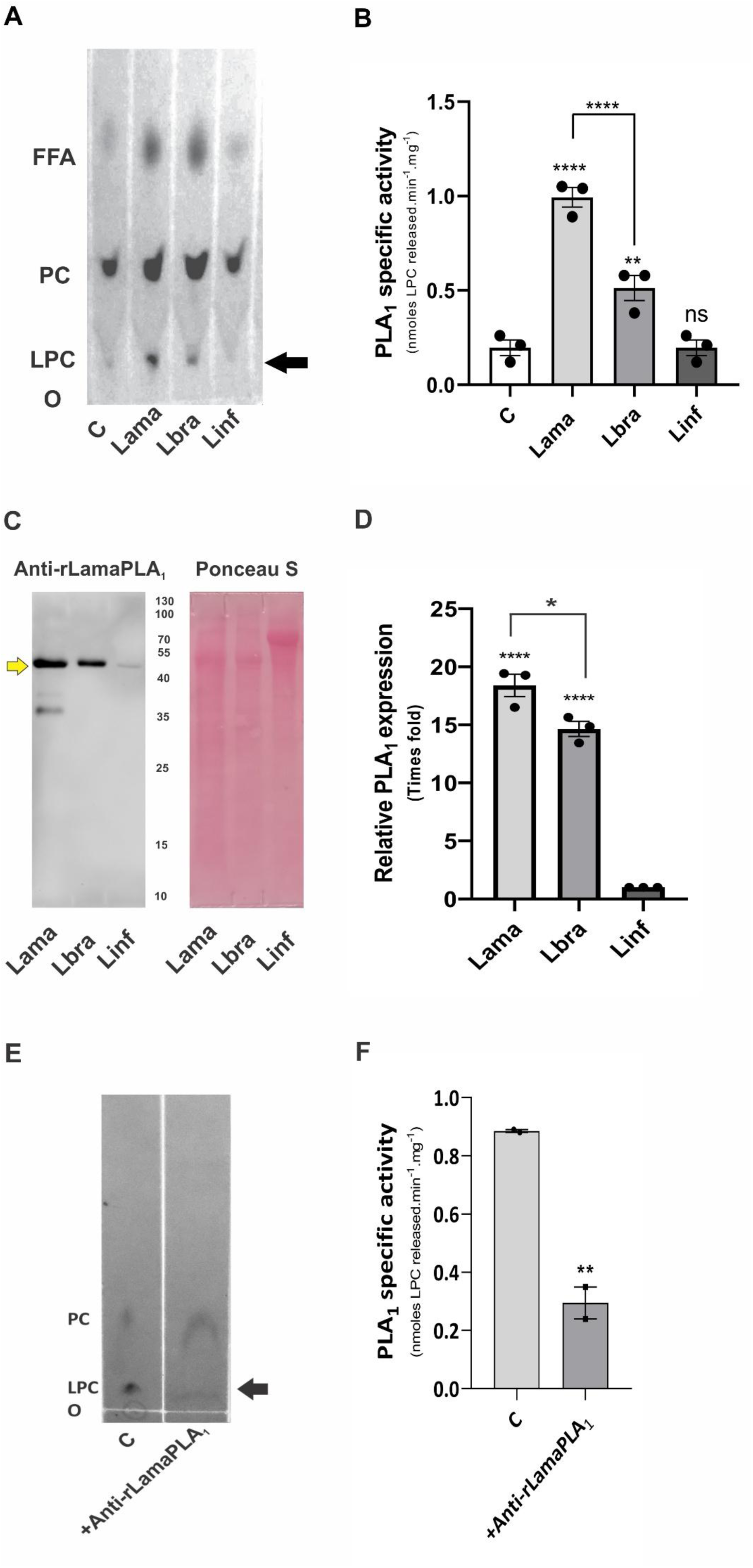
PLA_1_ expression and activity differ between *L. amazonensis* and *L. infantum* promastigotes. (A) PLA_1_ activity in parasite lysates of *L. amazonensis* (Lama), *L. braziliensis* (Lbra), and *L. infantum* (Linf) assessed by TLC. The substrate (control, **C**) was incubated under the same conditions, but without cell lysates. LPC generation (black arrow) is indicative of PLA_1_ activity in Lama and Lbra. FFA: free fatty acids, LPC: lysophosphatidylcholine, O: origin, PC: phosphatidylcholine. (B) PLA_1_ specific activity in parasite lysates. (C) Immunoblot analyses of PLA_1_ expression (∼50 kDa, yellow arrow) in Lama, Lbra, and Linf, using anti-rLamaPLA_1_ serum. (D) Relative PLA_1_ expression. Lama and Lbra PLA_1_ expressions were relativized to Linf for graphical analysis. (E) PLA_1_ activity in Lama pre-incubated with anti-rLamaPLA_1_ serum or normal mouse serum (control, C). Black arrow points at the LPC spot. (F) Inhibition of Lama PLA_1_ specific activity. Data are shown as mean ± SEM of at least two independent experiments with duplicate determinations. *p<0.05, **p<0.01, ****p<0.0001, ns: non-significant, one-way ANOVA.

In order to determine if the obtained levels of PLA_1_ activity were due to differential expression of the enzyme, immunoblot analyses were performed using a polyclonal serum against the recombinant *L. amazonensis* PLA_1_ (anti-rLamaPLA_1_), which was developed as detailed in Supplementary Fig. S1. Immunoblot shows major bands of approximately 50 kDa, with different intensities, in *L. amazonensis*, *L. braziliensis,* and *L. infantum* promastigotes lysates (Fig. 1C). The analysis of the different PLA_1_ protein bands was performed by densitometry, and it was found that the most intense band corresponded to *L. amazonensis*, followed by *L. braziliensis* and *L. infantum*. Protein levels of *L. amazonensis* PLA_1_ were approximately 1.3-fold higher than those of *L. braziliensis*, whereas *L. infantum* promastigotes showed a barely detectable signal (Fig. 1D). Although *L. infantum* promastigotes showed no detectable levels of PLA_1_ activity under our experimental conditions, immunoblot analyses confirmed the presence of PLA_1_ expression. These results suggest differences between PLA_1_ protein levels and enzymatic activity among the *Leishmania* species here analyzed.

We further characterized *L. amazonensis* PLA_1_ activity by testing if anti-rLamaPLA_1_ serum could inhibit this enzyme in promastigote lysates. PLA_1_ assays were performed after pre-incubating *L. amazonensis* promastigote lysates with anti-rLamaPLA_1_ serum or normal mouse serum (control). Fig. 1E shows that pre-incubation with anti-rLamaPLA_1_ serum decreased LPC generation with respect to the control. Densitometric analysis of TLC plates indicated that anti-rLamaPLA_1_ serum decreased 3-fold *L. amazonensis* PLA_1_ activity when compared to control (0.89 ± 0.01 and 0.30 ± 0.05 nmoles of LPC released.min^-1^.mg^-1^ protein, respectively) (Fig. 1F).

### 3.2. L. amazonensis PLA_1_ activity is modified by pH and divalent cations

Next, to determine if variations in pH and divalent cations could modify PLA_1_ activity, we tested *L. amazonensis* promastigote lysates under different conditions. Using a universal buffer adjusted to a range of pH values between 3.0 and 8.0, we determined that the highest PLA_1_ activity was achieved at pH 4.8 (Fig. 2A). Moreover, the addition of 2 mM Ca^2+^ or 2 mM Mg^2+^ significantly increased *L. amazonensis* PLA_1_ activity by approximately 50% and 30%, respectively (Fig. 2B).

**Fig. 2.**
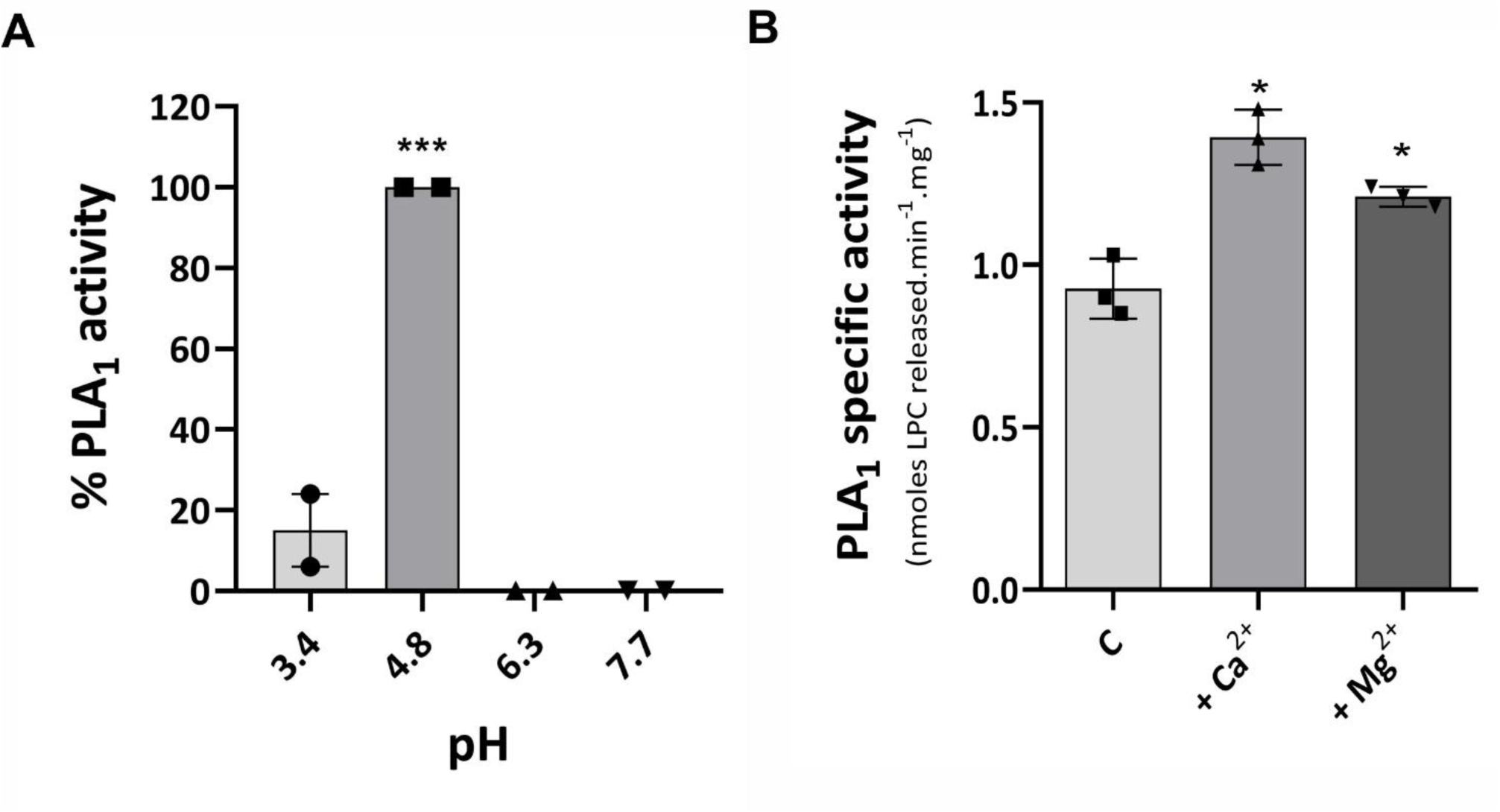
*L. amazonensis* PLA_1_ activity is modified by pH and divalent cations. (A) PLA_1_ activity was assayed in *L. amazonensis* promastigote lysates (Lama) at different pH values (3.0-8.0). (B) The effect of divalent cations on PLA_1_ activity was evaluated by incubating Lama with only the substrate (control, C), or supplemented with 2 mM Ca^2+^ (+Ca^2+^) or 2 mM Mg^2+^ (+Mg^2+^). Data are shown as mean ± SEM of at least two independent experiments with duplicate determinations. *p<0.05, ***p<0.001, one-way ANOVA.

### 3.3. Recombinant L. amazonensis PLA_1_ was successfully expressed and purified

A previous ortholog search in the TriTrypDB and NCBI databases for *Leishmania* PLA_1_ genes, based on the nucleotide sequence of *L. braziliensis* PLA_1_ (GenBank ACCN KJ957826), showed the existence of a putative lipase (class 3) gene in *L. amazonensis* (LAMA_000646500), among other species[14]. Therefore, we decided to carry out the recombinant expression of this gene to further characterize the corresponding protein. To this end, the partial gene sequence of putative *L. amazonensis* PLA_1_ (without the 30 amino acids from the predicted signal peptide) was commercially synthesized and cloned into the pET-28a (+) expression vector and expressed as described in the materials and methods section (2.5).

An amino acid sequence alignment was performed for PLA_1_ orthologs from *L. braziliensis* (GenBank ACCN AJT59458), *L. infantum* (GenBank ACCN LINJ_31_2540), the *L. amazonensis* full-length ortholog (TriTrypDB LAMA_000646500), and the recombinant *L. amazonensis* PLA_1_ (rLamaPLA_1_, GenBank ACCN WOA05485) [32]. A comparison of the deduced amino acid sequences of *L. amazonensis* with those of *L. braziliensis* and *L. infantum* indicated a 57% and 79% identity, respectively (Figure 3).

**Fig. 3.**
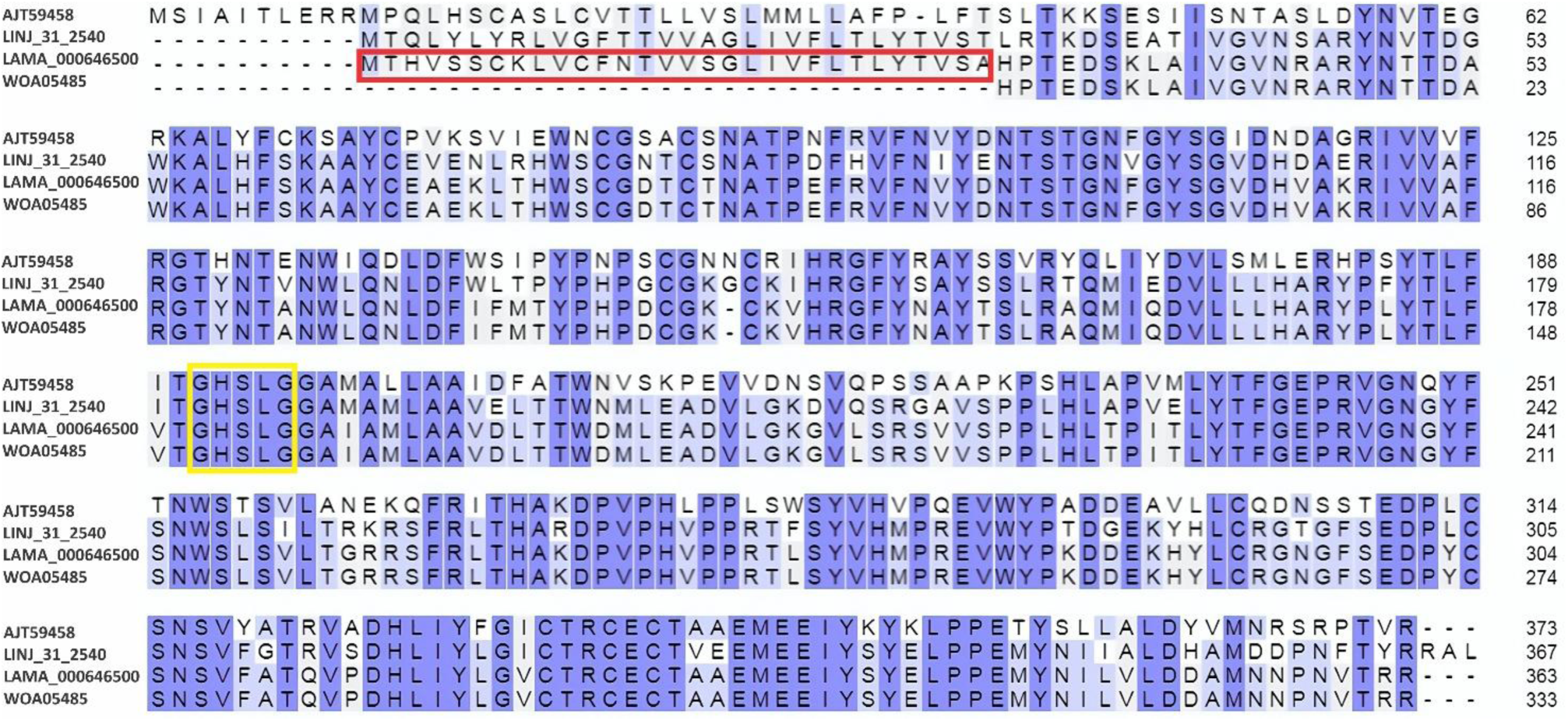
Alignment of the deduced amino acid sequences of PLA_1_ orthologs. Sequences from *L. braziliensis* (GenBank ACCN AJT59458), *L.infantum* (GenBank ACCN LINJ_31_2540), and the *L. amazonensis* full-length ortholog (TriTrypDB LAMA_000646500) that possesses the signal peptide (red box) are shown. The sequence for recombinant *L. amazonensis* PLA_1_ (rLamaPLA_1_, GenBank ACCN WOA05485) is also included. The lipase consensus pattern (GXSXG) is highlighted within the yellow box. Multiple sequence alignment was performed using the Align tool from the UniProt Database.

The expression of rLamaPLA_1_ in a prokaryotic system was successfully achieved, and a schematic representation of the process is shown in Supplementary Fig. S2. Since the His-tag recombinant protein accumulated as inclusion bodies (IBs), purification was performed under denaturing conditions using a Ni_++_-NTA resin. Fig. 4A shows the SDS-PAGE analysis of the purification steps, revealing a single band of approximately 40 kDa in lanes 5–7, corresponding to rLamaPLA_1_. The identity of this band was further confirmed by immunoblotting with anti-histidine antibodies (Fig. 4B). Altogether, successful *L. amazonensis* PLA_1_ recombinant expression was essential to obtain a specific anti-rLamaPLA_1_ serum (Supplementary Fig. S1), which in turn encouraged us to use it as a biological tool for exploring the subcellular localization of PLA_1_.

**Fig. 4.**
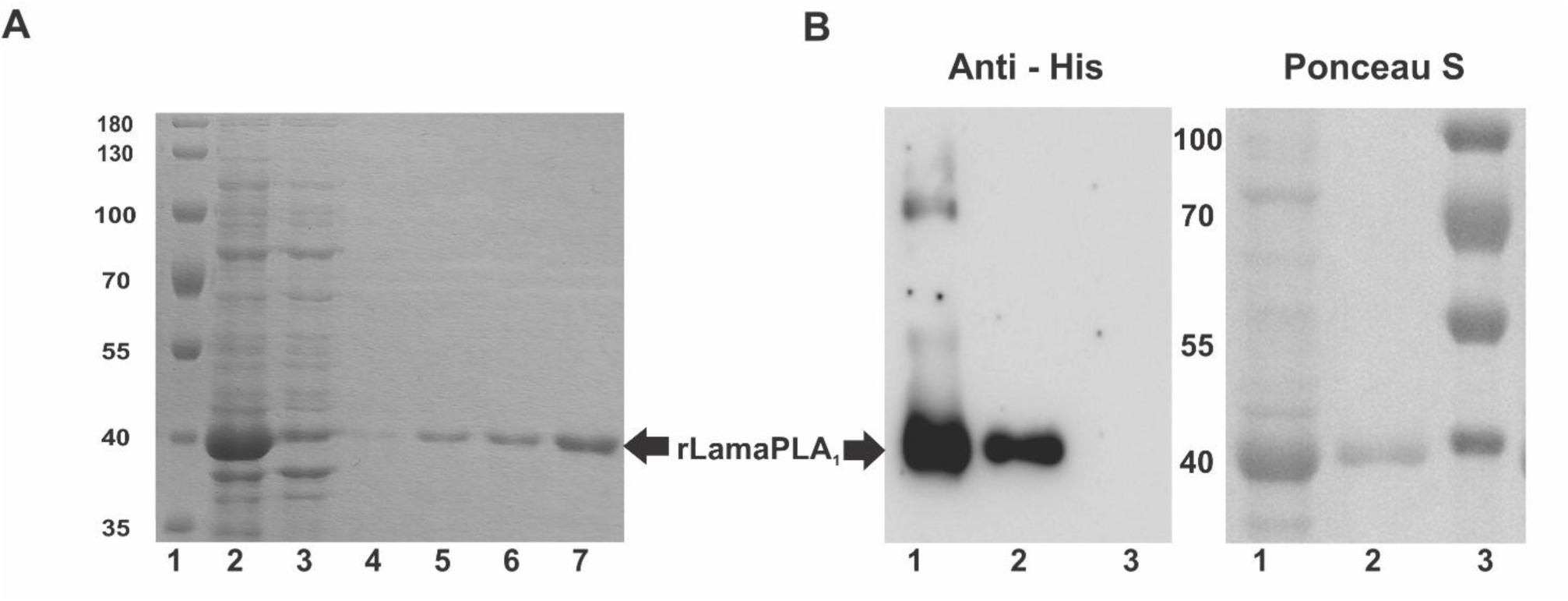
Expression and purification of recombinant *L. amazonensis* PLA_1_ (rLamaPLA_1_). (A) SDS-PAGE analysis of the purification steps. Black arrow points at the rLamaPLA_1_ band (∼40 kDa). Lanes: 1-molecular weight markers, 2-inclusion bodies, 3-flow through, 4-wash. 5, 6 and 7-eluates of increasing concentrations of imidazole (125, 250 and 400 mM, respectively). Images are representative of two independent experiments. (B) The identity of rLamaPLA_1_ was confirmed by immunoblot using an anti-histidine (anti-His) antibody, and bands were developed by chemiluminescence. As a loading control, Ponceau S staining was used. Lanes: 1-inclusion bodies, 2-purified rLamaPLA_1_, 3-molecular weight markers.

### 3.4. PLA_1_ and lipid droplets colocalize in Leishmania promastigotes

To investigate the subcellular localization of PLA_1_ within *L. amazonensis* and *L. infantum* promastigotes, we used anti-rLamaPLA_1_ serum. Lipid droplets (LD) were stained with the BODIPY probe, and colocalization with PLA_1_ was examined using confocal fluorescence microscopy. As shown in Fig. 5A, strong PLA_1_ immunoreactivity (red) was consistently detected in the cytoplasm and erratically in the flagella of *L. amazonensis* promastigotes. In addition, we analyzed the spatial relationship between PLA_1_ and LD. Merged images revealed regions of close association (yellow) between PLA_1_ and BODIPY-labeled LD, suggestive of colocalization (yellow arrows). Further semi-quantitative analysis from different fields validates a moderate to strong positive correlation between LD and PLA_1_ signals with a Pearson’s R value of 0.46 and 0.52 for each tested field (Fig. 5B). Moreover, orthogonal view from z-stacks of colocalization spots in images obtained from *L. amazonensis* indicate the presence of PLA_1_ in the outer layer of LD forming a cap (Fig. 5C), which is in agreement with the well characterized hydrophilic outer layer of LD and neutral hydrophobic core. Despite considerably reduced signal for both PLA_1_ and BODIPY-labeled LD in *L. infantum* promastigotes, PLA_1_ immunoreactivity (red) was predominantly found in the cytoplasm, as shown in Fig. 5D. This finding aligns with the lower expression levels found in parasite lysates compared to *L. amazonensis* (Figs. 1C and D). Similar to *L. amazonensis*, merged images revealed regions of PLA_1_ association with LD (yellow arrows), suggesting a similar pattern of distribution.

**Fig. 5.**
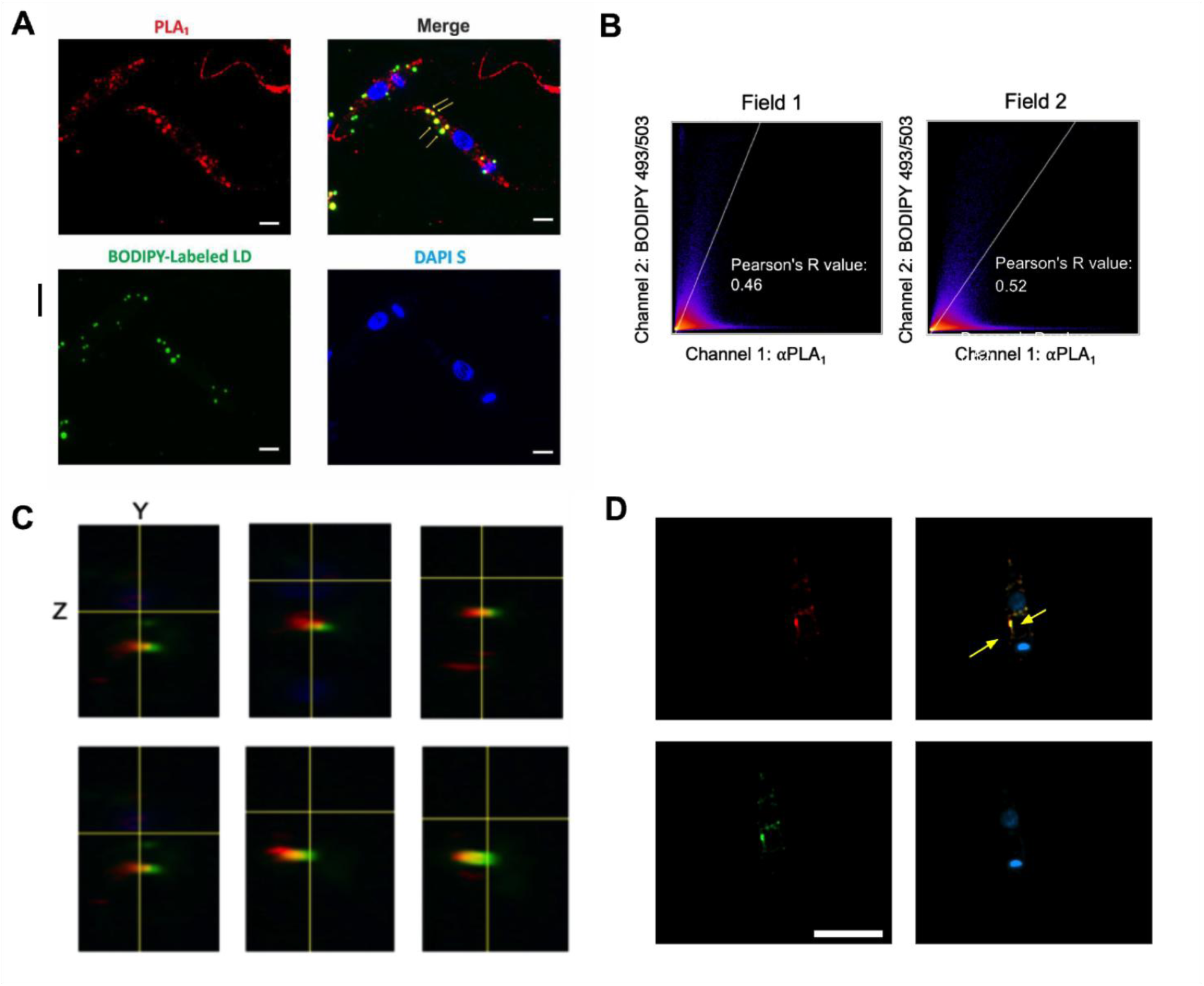
LD are sites for PLA_1_ expression. PLA_1_ and LD within *L. amazonensis* (A) visualized by confocal fluorescence microscopy and (B) semi-quantitative analysis of colocalization based on Pearson’s correlation. (C) Orthogonal view of Y and Z axis of LD colocalized with PLA_1_ staining and (D) PLA_1_ and LD within *L. infantum* promastigotes. PLA_1_ was detected using anti-rLamaPLA_1_ serum (red), the LD were stained with BODIPY^TM^ 493/503 (green), nuclei and kinetoplasts were identified with DAPI staining (blue). The merged image indicates the sites of colocalization between PLA_1_ and LD (yellow arrows). Figure shows representative zoomed images from two independent experiments. Scale bar: 2 μm (A) and 5 μm (D).

The recent release of the TrypTag resource [27] provided the opportunity to explore the subcellular localization of the homologous *T. brucei* PLA_1_ (TbPLA_1_). As shown in Supplementary Fig. S3, TbPLA_1_ (green) exhibited a heterogeneous distribution, with signals detected in the cytoplasm, endocytic compartments, and the flagellum. These patterns suggest that PLA_1_ occupies multiple subcellular niches, possibly reflecting distinct functional roles. *In silico* copredictions parallel our experimental observations in *L. amazonensis* and *L. infantum*, where PLA_1_ consistently localized to the cytoplasm and was associated with LD. In *L. amazonensis*, additional localization to the flagellum was observed. Together, these results support a consistent localization profile for PLA_1_ across the trypanosomatids tested, strengthening the biological relevance of the experimental data.

## 4. Discussion

Following our previous findings regarding *L. braziliensis* PLA_1_, we performed a comparative analysis of PLA enzymes in *L. amazonensis* and *L. infantum*, focusing on their expression, activity, and subcellular localization. We here report the presence of phosphatidylcholine-specific PLA_1_ activity in *L. amazonensis* promastigotes. This finding significantly expands the understanding of PLA activities in this species, which was previously restricted to descriptions of PLA_2_ activity [15,16,30,31].

*L. amazonensis* exhibited higher PLA_1_ protein expression than *L. braziliensis,* and accordingly, significant differences were observed in enzymatic activity between both species. In contrast, the absence of PLA_1_ activity and barely detectable PLA_1_ expression in *L. infantum* could be attributed to species-specific factors that could modulate them as previously described for other trypanosomatids. When comparing PLA_1_ specific activity levels from the vector’s parasite stages of pathogenic trypanosomes, we observed that *Leishmania* promastigotes, here analyzed, had the lowest values with respect to those already reported for *T. cruzi* epimastigotes and *T. brucei* procyclic trypomastigotes [14,29,33].

On the other hand, greater discrepancies between *L. amazonensis* and *L. infantum* regarding activity levels could be related not only to protein expression levels, but also to post-translational regulatory mechanisms, such as protein stability, activation, or inhibition, that may influence PLA_1_ activity. Besides, these activity levels could vary according to the substrate used, thus depending on the fatty acid composition or the polar head of the phospholipids [33]. Both *L. amazonensis* and *L. braziliensis* exhibited optimal activity under acidic pH, while showing a slight divergence in optimal pH values (4.8 and 6.4, respectively). The enzymatic activity was significantly enhanced in the presence of Ca_2+_ and Mg^2+^ ions. This cationic regulation is a distinct feature, as other PLAs activities reported in various trypanosomatids have been described as being independent of divalent cation supplementation [14]. This dependency suggests that *L. amazonensis* PLA_1_ is regulated by these divalent cations, through a binding site or activation mechanism. Moreover, the enzymatic activity within *L. amazonensis* promastigote lysates was inhibited *in vitro* by the polyclonal serum specific for *L. amazonensis* PLA_1_. These results are in agreement with previous findings from our group, which demonstrate that specific antibodies raised against this enzyme were capable of neutralizing *T. cruzi* PLA_1_ activity *in vitro* and *in vivo* [29]. Further studies are required to elucidate the factors governing PLA_1_ enzymatic regulation in these species.

A valid strategy to study genes and its associated protein functions is to perform recombinant protein expression. In this regard, our pioneering work provided the first molecular and functional characterization of a PLA_1_ in this genus. Previously, we successfully cloned and expressed the PLA_1_ gene from *L. braziliensis*, confirming its catalytic activity and establishing the foundation for the subsequent phylogenetic analysis of this enzyme across several *Leishmania* species [14]. In the present work, we cloned and expressed the *L. amazonensis* PLA_1_ gene in *E. coli* BL21-pLys. Although the recombinant protein accumulated primarily as IBs, this fact was strategically leveraged for purification, allowing the isolation of the target protein to a high degree of purity, following denaturation and refolding protocols. This high-quality recombinant protein served as an excellent immunogen for the generation of anti-rLamaPLA_1_ serum, proving to be a valuable and essential tool for biological assays, since commercial anti-PLA_1_ antibodies were raised against human and rat PLAs and did not recognize the enzyme in trypanosomatids. In this sense, this polyclonal specific serum allowed the detection of PLA_1_ protein expression not only in *L. amazonensis* but also in *L. infantum* and *L. braziliensis* promastigote lysates, in accordance with the high degree of identity in amino acid sequences among species.

A major contribution of the present study is the first description of the subcellular localization of a PLA_1_ within the *Leishmania* genus, achieved through confocal fluorescence microscopy. Our immunolocalization data revealed that the PLA_1_ enzyme exhibits a distinct pattern across the species investigated. Specifically, in both *L. amazonensis* and *L. infantum* promastigotes, the enzyme was clearly localized to the cytoplasm. Furthermore, in *L. amazonensis*, we observed an additional and striking localization of PLA_1_ within the flagellum. This dual localization in *L. amazonensis* is particularly noteworthy, suggesting a potential role for the flagellar PLA_1_ in lipid turnover associated with the flagellar membrane or motility, warranting further investigation into the mechanism of its targeted transport. Further refinement of the PLA_1_ localization data revealed an unprecedented association between this enzyme and LD within the *Leishmania* parasite. Confocal microscopy demonstrated significant colocalization of the PLA_1_ signal with BODIPY labeled-LD across both *L. amazonensis* and *L. infantum* promastigotes. This finding is of considerable interest, as it represents the first evidence linking any PLA_1_ to LD in the *Leishmania* genus.

*In silico* analyses using the TrypTag resource [27] predicted a similar complex localization (cytoplasmic, flagellar, or endocytic) for the homologous PLA_1_ in *T. brucei*, providing strong support for the experimental findings observed in *L. amazonensis* and *L. infantum*. This parallel, *in silico* and experimental patterns, suggest that PLA_1_ occupies multiple subcellular niches across trypanosomatids, likely reflecting distinct functional roles. Together, these results support a conserved localization profile for PLA_1_, strengthening the biological relevance of our data.

The localization of PLA_1_ within *Leishmania* LD is of substantial biological significance when evaluated against the background of eicosanoid metabolism in trypanosomatids. Previous works have established that parasite LD are crucial intracellular sites for eicosanoid synthesis, with LD induction correlating with the expression of prostaglandin F_2α_ (PGF_2α_) synthase in *L. infantum chagasi* [34] and serving as sites for PGE2 synthesis in *T. cruzi* upon AA stimulation [35]. Given that eicosanoid pathways are initiated by the release of fatty acid precursors, most notably AA from phospholipids, our demonstration of a PLA_1_ in direct spatial association with LD places a key regulatory enzyme right at the metabolic nexus. This localization strongly suggests that PLA_1_ plays an integral role in modulating the availability of these lipid substrates, thereby influencing the synthesis of immune-modulating molecules essential for parasite pathogenesis and survival.

In conclusion, this study constitutes the first report identifying PLA_1_ in *L. amazonensis* and *L. infantum*, while providing a detailed characterization of their enzymatic activity, expression profiles, and subcellular localization within the cytoplasm associated with LD. These findings are particularly significant given that PLA_1_s are established virulence factors in other trypanosomatids. Furthermore, considering that *Leishmania* has evolved sophisticated mechanisms to modulate the host immune response, ensuring its survival and persistence, our results highlight the need for future research focused on the role of this enzyme in LD dynamics and eicosanoids generation. Understanding how PLA_1_-derived lipid mediators contribute to this immune subversion may be crucial for unraveling parasite pathogenesis and identifying potential therapeutic targets.

## Data availability statement

All data generated or analyzed during this study are included in the article/supplementary material, further inquiries can be directed to the corresponding author.

## Acknowledgements

We thank Drs. Victoria Fragueiro Frías and Vanesa Negri (ANLIS-Instituto Nacional de Parasitología “Dr. Mario Fatala Chabén”, Buenos Aires, Argentina) for providing the *Leishmania (L.) amazonensis* IFLA/BR/67/PH8 reference strain, and Prof. Sandra V. Verstraeten (Instituto de Química y Fisicoquímica Biológicas - CONICET/UBA, Buenos Aires, Argentina) for assisting us in the use of the fluorometer. We are also grateful to Dr. Korenaga Masataka and Dr. Hashiguchi Yoshitaka for their encouragement throughout the research on leishmaniases. Maintenance of the *Leishmania (L.) infantum* MHOM/MA/67/ITMAP263 reference strain for this work was conducted as part of Juan José Lauthier’s Ph.D. scholarship at Kochi Medical School, Kochi University, Japan, funded by the Japan Ministry of Education, Culture, Sports, Science and Technology (MEXT), and partially supported by the Institute of Tropical Medicine (NEKKEN), Nagasaki University, Japan.

## Authorship contribution statement

SAL, TSSV, SNT, GG and MLB conceived the study, designed the experiments, provided the information, analyzed the data, visualization and wrote the first draft. MAR, ASA, JJL, PTB, GG and MLB contributed funding acquisition, reagents, materials, and analysis tools. FSPD, MGR and AEB reviewed and critically revised and improved the paper. All authors have read and agreed to the published version of the manuscript.

