## Supplementary Figures for "Identification, expression and subcellular localization of *Leishmania amazonensis* and *Leishmania infantum* Phospholipases A_1_"

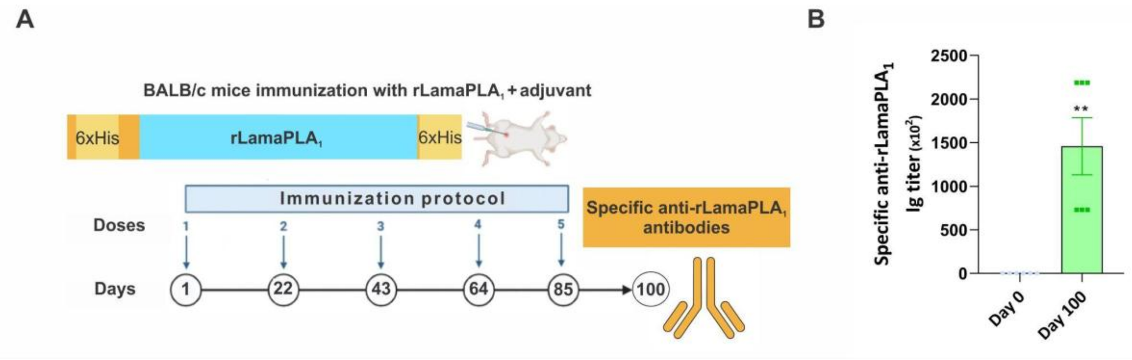


Supplementary Fig. S1. Generation of polyclonal specific anti-rLamaPLA$_1$ antibodies. (A) Mice immunization protocol (*n* = 6) with rLamaPLA$_1$ (Image created using BioRender). (B) Specific anti-rLamaPLA$_1$ immunoglobulin (Ig) titer assessment in mice sera. Data are shown as mean ± SEM, **$p<0.01$, unpaired Student's t-test.

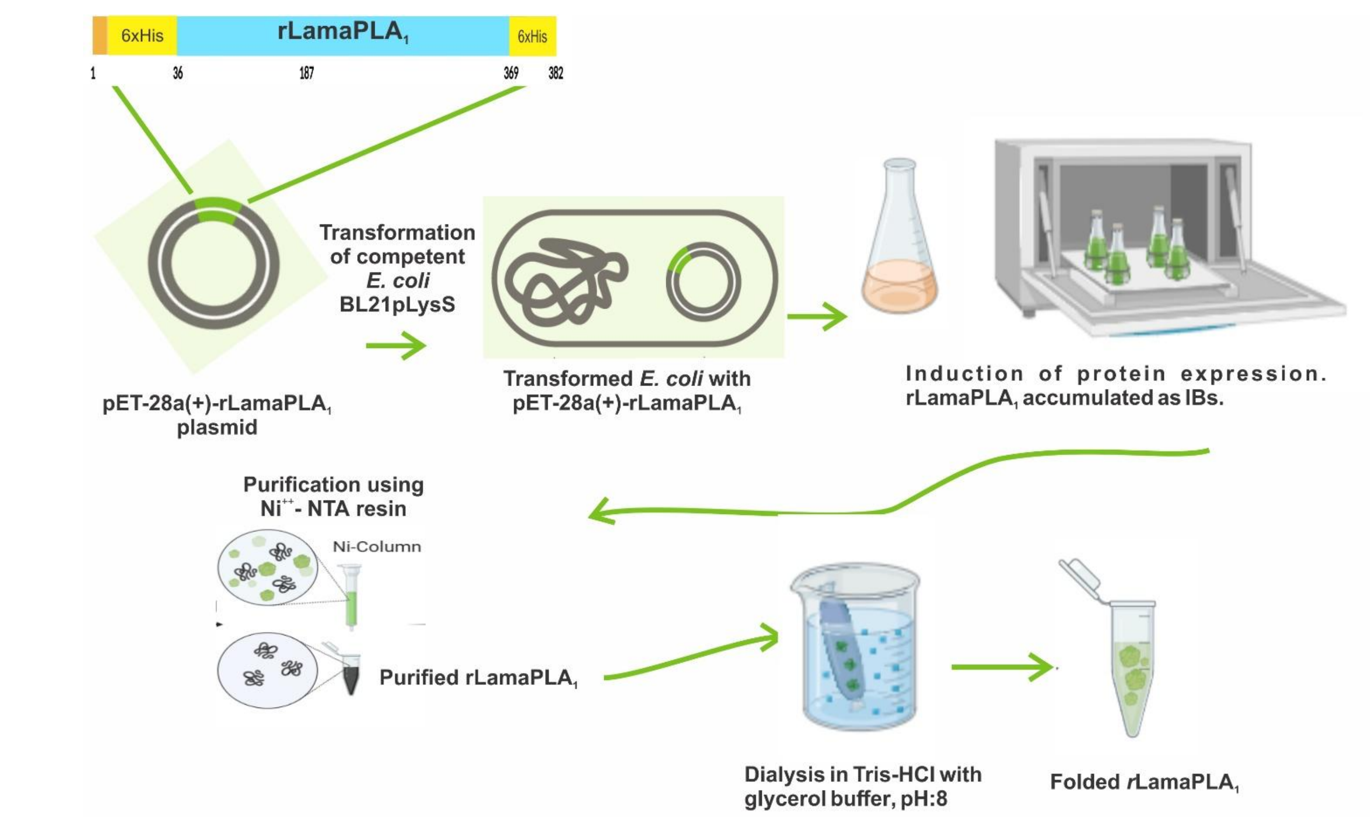


Supplementary Fig. S2. Flux diagram of recombinant *L. amazonensis* PLA$_1$ (rLamaPLA$_1$, GenBank WOA05485) expression and purification. Briefly, the pET-28a(+)-rLamaPLA$_1$ construct was used to transform competent *E. coli* BL21pLysS cells. Transformed bacteria were induced for protein expression for 20 h at 24°C. Since the rLamaPLA$_1$ accumulated as inclusion bodies (IBs), it was purified under denaturing conditions using $Ni^{++}$-NTA resin and further refolded *in-vitro* by dialysis. 6xHis: His tags (yellow boxes). Image created using BioRender.

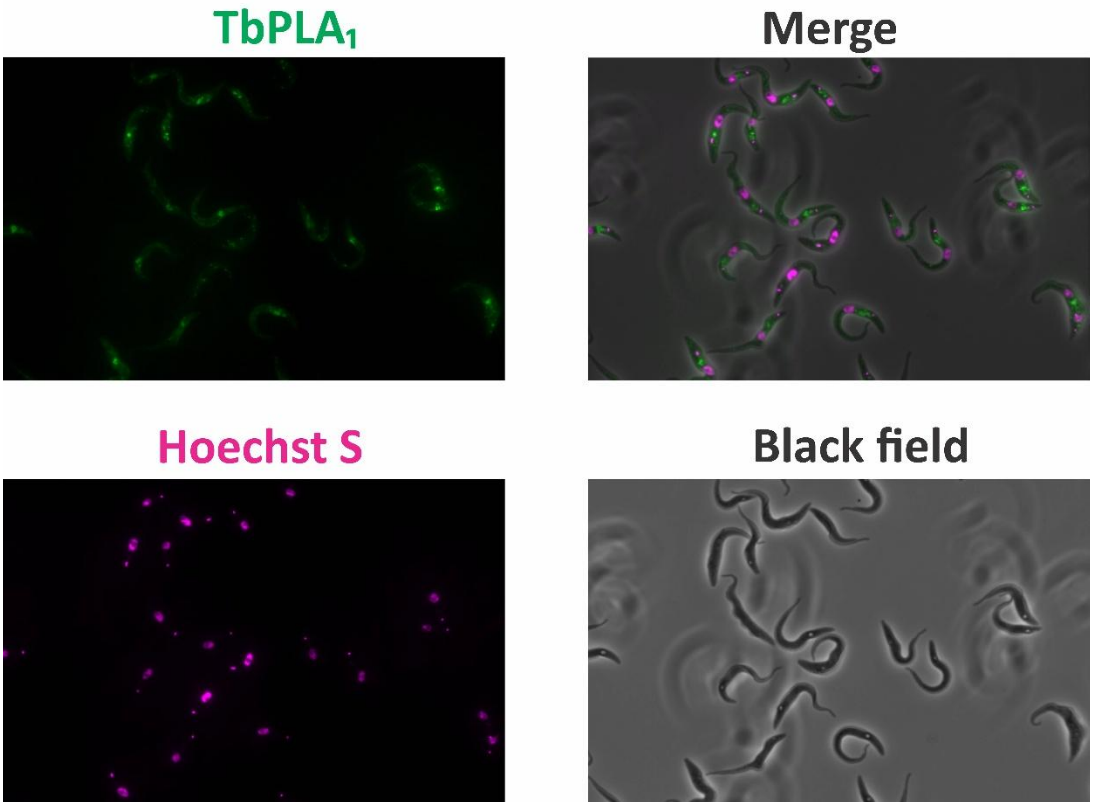


Supplementary Fig. S3. *In silico* analysis of *T. brucei* PLA$_1$ (TbPLA$_1$). The TrypTag resource was used to explore the subcellular localization of TbPLA$_1$ (Tb927.8.7440, TREU927 reference strain). Hoechst 33342 (violet) was used for staining nuclei and kinetoplasts. The figure shows representative zoomed epifluorescence images.
